# AXL is a key factor for cell plasticity and promotes metastasis in pancreatic cancer

**DOI:** 10.1101/2020.07.06.190363

**Authors:** Wenting Du, Huocong Huang, Zhaoning Wang, Jason E. Toombs, Natalie Z. Phinney, Yuqing Zhang, Muhammad S. Beg, Thomas M. Wilkie, James B. Lorens, Rolf A. Brekken

## Abstract

Pancreatic ductal adenocarcinoma (PDA), a leading cause of cancer-related death in the US, has a high metastatic rate and is associated with persistent immune suppression. AXL, a member of the TAM (TYRO3, AXL, MERTK) receptor tyrosine kinase family, has been identified as a driver of metastasis and immune suppression in multiple cancer types. Here we use single cell RNA sequencing to reveal that AXL is expressed highly in tumor cells that have a mesenchymal-like phenotype and that AXL expression correlates with classic markers of mesenchymal tumor cells. We demonstrate that *AXL-deficiency* extends survival, reduces primary and metastatic burden and enhances sensitivity to gemcitabine in an autochthonous model of PDA. PDA in *AXL-deficient* mice displayed a more differentiated histology, higher nucleoside transporter expression and a more active immune microenvironment compared to PDA in *wild-type* mice. Finally, we demonstrate that AXL-positive mesenchymal tumor cells are critical for PDA progression and metastasis, emphasizing the potential of AXL as a therapeutic target for PDA patients.

## Introduction

Pancreatic ductal adenocarcinoma (PDA) is an aggressive cancer that has seen limited improvement in overall survival in the past 30 years (1). More than 80% of the patients are diagnosed at a late stage and as a result are not candidates for curative intent surgery (1). Thus, systemic chemotherapy is the mainstay of treatment. Current regimens, (5-fluorouracil or gemcitabine-based) offer only incremental prolongation of survival. Tumor cell intrinsic and acquired resistance to standard therapy underlies the poor response to systemic therapy (2). Tumor cell epithelial plasticity is a major contributor to PDA resistance to therapy.

There is a strong association between mesenchymal cancer cells and therapy resistance in multiple types of cancers (3–7). Tumor cells with a mesenchymal-like phenotype have a distinct gene expression signature, which often includes AXL (4, 8). AXL is a member of the TAM (TYRO3, AXL, MERTK) receptor tyrosine kinase family that binds to growth arrest-specific gene 6 (GAS6). AXL via GAS6 activation and via non-ligand-dependent mechanisms (9) mediates basic cell biological processes including cell migration and survival (10). Consistent with gene expression profiling studies, AXL expression is associated with therapy-resistance and metastasis in a variety of cancers, including NSCLC, breast cancer and PDA (11–20). AXL is also expressed by immune cells where it functions in efferocytosis, the clearance of apoptotic cells. This occurs in a GAS6-dependent manner. GAS6 contains a γ-carboxylation (Gla) domain that interacts with phosphatidylserine exposed on apoptotic cells. Thus, GAS6 serves as a bridging protein that results in the engulfment of apoptotic cells via AXL (or MERTK) activation (21, 22). Activation of AXL on immune cells in this manner also has immune regulatory effects (23–27).

In PDA, high AXL expression is detected in ~70% of patients and is associated with distant metastasis and worse survival (28, 29). Our lab previously investigated different pharmacologic strategies to inhibit AXL activity in PDA (30, 31). We found that inhibition of vitamin K-dependent γ-carboxylation of Gla domain of GAS6 with warfarin suppressed metastasis of preclinical models of PDA in a tumor cell AXL-dependent manner (31), that warfarin use was associated with improved outcome in PDA patients (32) and that warfarin use is associated with reduced incidence of multiple cancer types, including pancreatic cancer (33). We also demonstrated the efficacy of a selective small molecular inhibitor of AXL, bemcentinib (BGB324) in combination with gemcitabine in multiple mouse models of PDA. We found that BGB324 drove epithelial differentiation, induced an immune-stimulatory environment and improved gemcitabine efficacy in vivo (30). However, the distribution of AXL expression on different cells within PDA microenvironment and their contribution to PDA progression have not been investigated. Here we use single cell RNA sequencing (scRNA-Seq) to identify AXL-positive cell populations in genetically engineered mouse models (GEMMs) of PDA. We demonstrate the effect of AXL deletion on these AXL-positive populations, as well as PDA progression, metastasis and chemoresistance. Lastly, we show that AXL-positive tumor cells are critical for PDA progression. Our findings bolster the preclinical rationale for pharmacologic inhibition of AXL in PDA patients.

## Results

### AXL is expressed in mesenchymal PDA tumor cells

We and others have shown that multiple cell types in the tumor microenvironment can express AXL (9, 10). To determine which cell populations in GEMMs of PDA are AXL positive, we analyzed scRNA-Seq data (34) with tumors harvested from early and late stage of *KIC* (*Kras*^*LSL-G12D*/+^, *Ink4a/Arf*^*lox/lox*^, *Ptf1a*^*Cre*/+^) and late stage *KPfC* (*Kras*^*LSL-G12D*/+^, *Trp53*^*lox/lox*^, *Pdx*^*Cre*/+^) mice. tSNE plots of *Axl* expression for each cell population in *KIC* and *KPfC* tumors are shown in **Supplementary Figure 1**. *Axl* expression is highlighted in red for each model. Violin plots show that *Axl* is expressed highly in advanced *KIC* (**Fig. 1A**) and is expressed by mesenchymal cancer cells, fibroblasts and macrophages in each GEMM (**Fig. 1B**). To confirm the scRNA-seq data, tumor tissues from *Axl*^*LacZ*/+^ *KIC* mice were evaluated for co-localization of β-galactosidase (identifying AXL expression, red) and PDA cell marker SOX9, fibroblast marker PDGFRα and αSMA, macrophage marker F4/80, epithelial marker E-cadherin and mesenchymal marker Vimentin. Double-positive cells were found using each of the markers except E-cadherin (**Fig. 1C**).

**Figure 1.**
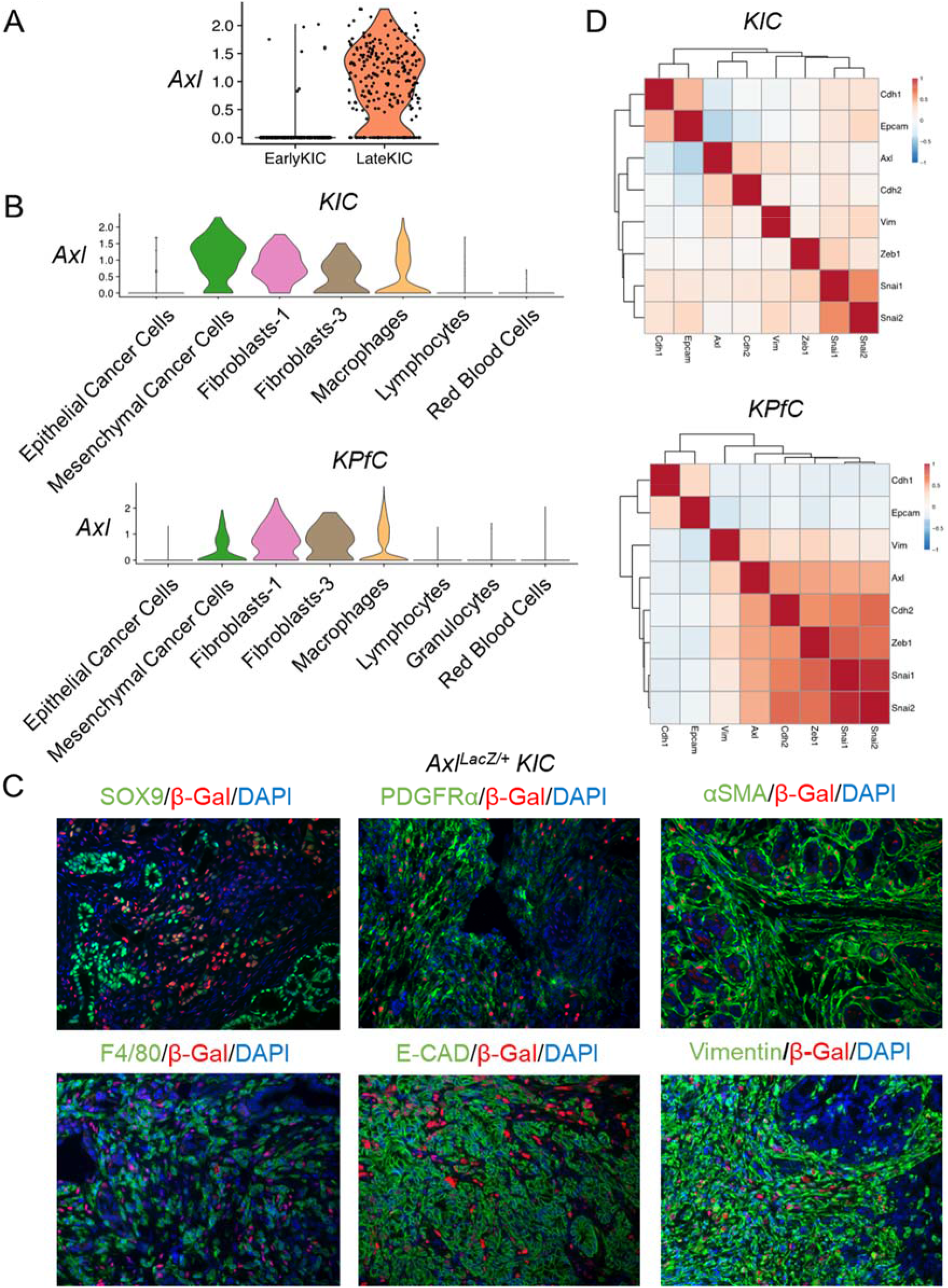
AXL is expressed in mesenchymal PDA tumor cells. **A)** Single cell RNA sequencing was performed with tumors harvested from early stage and late stage of *KIC* mice. *Axl* expression is shown by violin plot. **B)** Single cell RNA sequencing was performed with tumors harvested from *KIC* (*Kras*^*LSL-G12D*/+^, *Ink4a/Arf*^*lox/lox*^, *Ptf1a*^*Cre*/+^) or *KPfC* (*Kras*^*LSL-G12D*^, *P53*^*lox/lox*^, *PDX*^*Cre*/+^) mice. *Axl* expression in different populations is shown by violin plot for each mouse model. **C)** Tumor tissues from *Axl*^*LacZ*/+^ *KIC* mice were evaluated for co-localization of β-galactosidase and indicated targets. β-galactosidase staining is used to identify the expression of AXL. **D)** Correlation plots with *Axl* and EMT genes in *KIC* and *KPfC* models were generated by analyzing the EMT gene expression within the cancer cell clusters from each model. A positive correlation is highlighted in red and negative correlation is highlighted in blue.

Gene clusters from AXL-positive cell populations were subjected to pathway and Gene Ontology (GO) analysis. The most highly expressed genes in AXL-positive cancer cells were associated with focal adhesion pathway, MAPK cascade, PI3K/AKT signaling, ECM-receptor interaction, and NF-κB signaling as well as functions that were regulated by these signaling pathways including wound healing, cell migration, cell adhesion, cell senescence, cell growth, epithelial to mesenchymal transition (EMT), and response to drug (**Supplementary Fig. 2**). *Axl* expression correlated with mesenchymal markers and negatively correlated with epithelial marker expression in cancer cell clusters in each GEMM (**Fig. 1D**), highlighting that AXL is associated with EMT in PDA tumor cells. In AXL-positive fibroblasts, the most highly expressed genes were involved in fibril organization (p=4.07×10^−18^), extracellular matrix organization (p=2.55×10^−15^) and collagen formation (p=8.96×10^−13^). Whereas in AXL-positive macrophages, the most highly expressed genes were the associated with antigen processing and presentation (p=1.05×10^−11^).

### AXL deficiency prolongs survival, inhibits metastasis and EMT, and improves gemcitabine efficacy in a GEMM of PDA

To demonstrate the function of AXL in PDA progression, *Axl*^*LacZ/LacZ*^ mice (*lacZ* gene was inserted into the *Axl* locus) were crossed with *KIC* (*Kras*^*LSL-G12D*/+^, *Ink4a/Arf*^*lox/lox*^, *Ptf1a*^*Cre*/+^) mice (35). Global depletion of AXL in *KIC* mice (n=26) led to a 10-day longer median survival (68 days, p<0.0001) compared to *wild-type* (*WT*) *KIC* (58 days, n=21) animals (**Fig. 2A**). *Axl*^*LacZ/LacZ*^ *KIC* mice had smaller primary tumors and fewer liver micrometastases (**Fig. 2A**). Further analysis of tumor tissue from *WT* and *Axl*^*LacZ/LacZ*^ *KIC* mice showed that global AXL depletion did not affect cell proliferation (Ki67) or cell apoptosis (cleaved-caspase3, CC3, **Supplementary Fig. 3**). However, hematoxylin and eosin (H&E) and EMT marker staining showed that AXL deficiency resulted in a less advanced, and more differentiated histology (increased E-cadherin and decreased Vimentin) (**Fig. 2B**). Global depletion of AXL also led to reduced αSMA expression by cancer associated fibroblasts (CAFs) indicative of reduced myCAFs (34) in the absence of AXL (**Fig. 2B**). In contrast, PDGFRα and CDH11, other markers of CAFs, and CD31, a marker of microvessel density, were not affected significantly by AXL deficiency (**Supplementary Fig. 3**).

**Figure 2.**
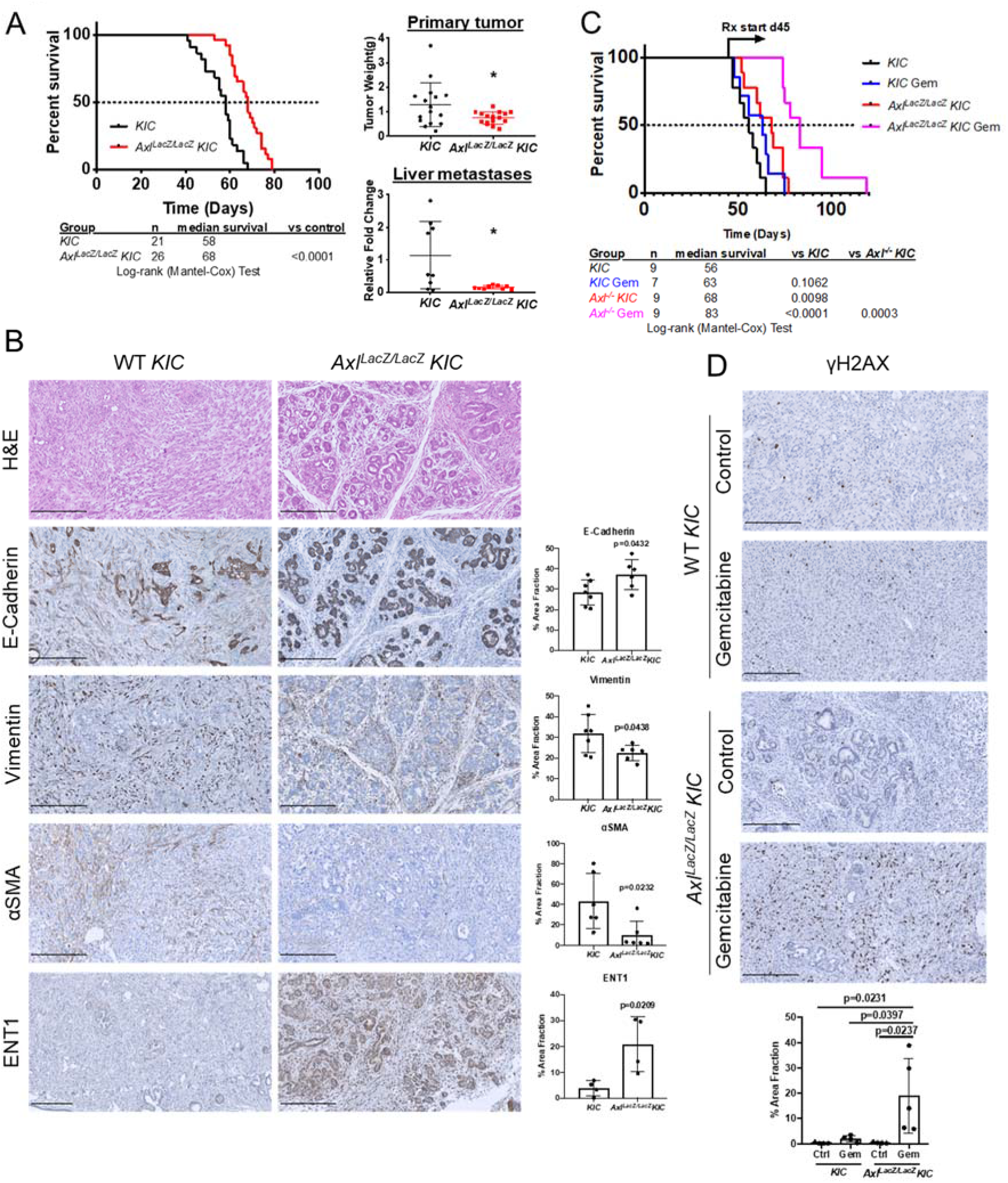
AXL deficiency prolongs survival, inhibits metastasis and EMT, and improves gemcitabine efficacy in a GEMM of PDA. **A)** *KIC* (n=21) and *Axl*^*LacZ/LacZ*^ *KIC* (n=26) mice were enrolled in a survival study. Primary tumor weight and liver metastases were measured at the end of the survival study. Liver metastases were determined by detecting recombined *Ink4a/Arf* in liver tissues. Data are displayed as mean ± SD. *P < 0.05; by t-test. **B)** Tumor tissues from *KIC* and *Axl*^*LacZ/LacZ*^ *KIC* mice were evaluated for indicated targets. Images were taken by Nanozoomer to cover the whole tissue and analyzed using ImageJ software. quantification of % area fraction is shown. Each data point is the average of one sample. Data are displayed as mean ± SD. P values were generated by t-test. Scale bar, 250 μM. **C)** *KIC* and *Axl*^*LacZ/LacZ*^ *KIC* mice were enrolled in a survival study and randomized to vehicle and gemcitabine (Gem, 25 mg/kg ip twice a week). Therapy was initiated on day 45 and maintained until sacrifice. **D)** Tumor tissues from **(C)** were evaluated for γH2AX. Images were taken by Nanozoomer to cover the whole tissue and analyzed using ImageJ software. Quantification of % area fraction is shown. Each data point is the average of one sample. Data are displayed as mean ± SD. P values were generated by ANOVA. Scale bar, 250 μM.

EMT is associated with chemoresistance (5, 6). EMT can reduce the expression of nucleoside transporters, which limits the entry of chemotherapeutics into cancer cells (5). Given the more differentiated phenotype of *AXL-deficient KIC* tumors we stained *WT* and *AXL-deficient* tumors for expression of equilibrative nucleoside transporter 1 (ENT1). Consistent with expectations, we found a dramatic increase of ENT1 protein expression in *Axl*^*LacZ/LacZ*^ *KIC* compared to *WT KIC* animals (p=0.0209, **Fig. 2B**). Based on this result, we hypothesized that *Axl*^*LacZ/LacZ*^ *KIC* mice would be more sensitive to chemotherapy (gemcitabine). To test this, *WT KIC* and *Axl*^*LacZ/LacZ*^ *KIC* were treated with vehicle (*KIC* n=9 and *Axl*^*LacZ/LacZ*^ *KIC* n=9) or gemcitabine (25 mg/kg, twice a week; *KIC* Gem, n=7; *Axl*^*LacZ/LacZ*^ Gem, n=9) until animals were moribund. Therapy was initiated at 45 days of age. *WT KIC* mice that received vehicle had a median survival of 56 days, while treatment with gemcitabine only modestly extended the median survival by 12.5% to 63 days (p=0.1062 vs. *KIC*). Similar to **Figure 2A**, *Axl*^*LacZ/LacZ*^ *KIC* treated with vehicle had a relatively longer median survival (68 days) compared to vehicle treated *WT KIC* (p=0.0098 vs. *KIC*). However, gemcitabine treatment in *Axl*^*LacZ/LacZ*^ *KIC* significantly improved survival by 22.1% to a median of 83 days (p=0.0003 vs. *Axl*^*LacZ/LacZ*^ *KIC*; **Fig. 2C**). Consistent with the survival result, γH2AX staining in tumor tissues demonstrated that gemcitabine-induced DNA double-strand break were potently promoted in *Axl*^*LacZ/LacZ*^ *KIC* animals (**Fig. 2D**).

### AXL deficiency results in a more active immune microenvironment

To investigate how AXL contributes to the immune landscape of PDA, RNA was isolated from *KIC* (n=2) and *Axl*^*LacZ/LacZ*^ *KIC* (n=3) tumors and was analyzed using Mouse PanCancer Immune Profiling Panel (NanoString Technologies). Upregulated (Red) and downregulated (Blue) gene programs/functions are shown in **Figure 3A**. Most immune cell functions as defined by NanoString were upregulated in *Axl*^*LacZ/LacZ*^ *KIC*, indicating a more active immune microenvironment.

**Figure 3.**
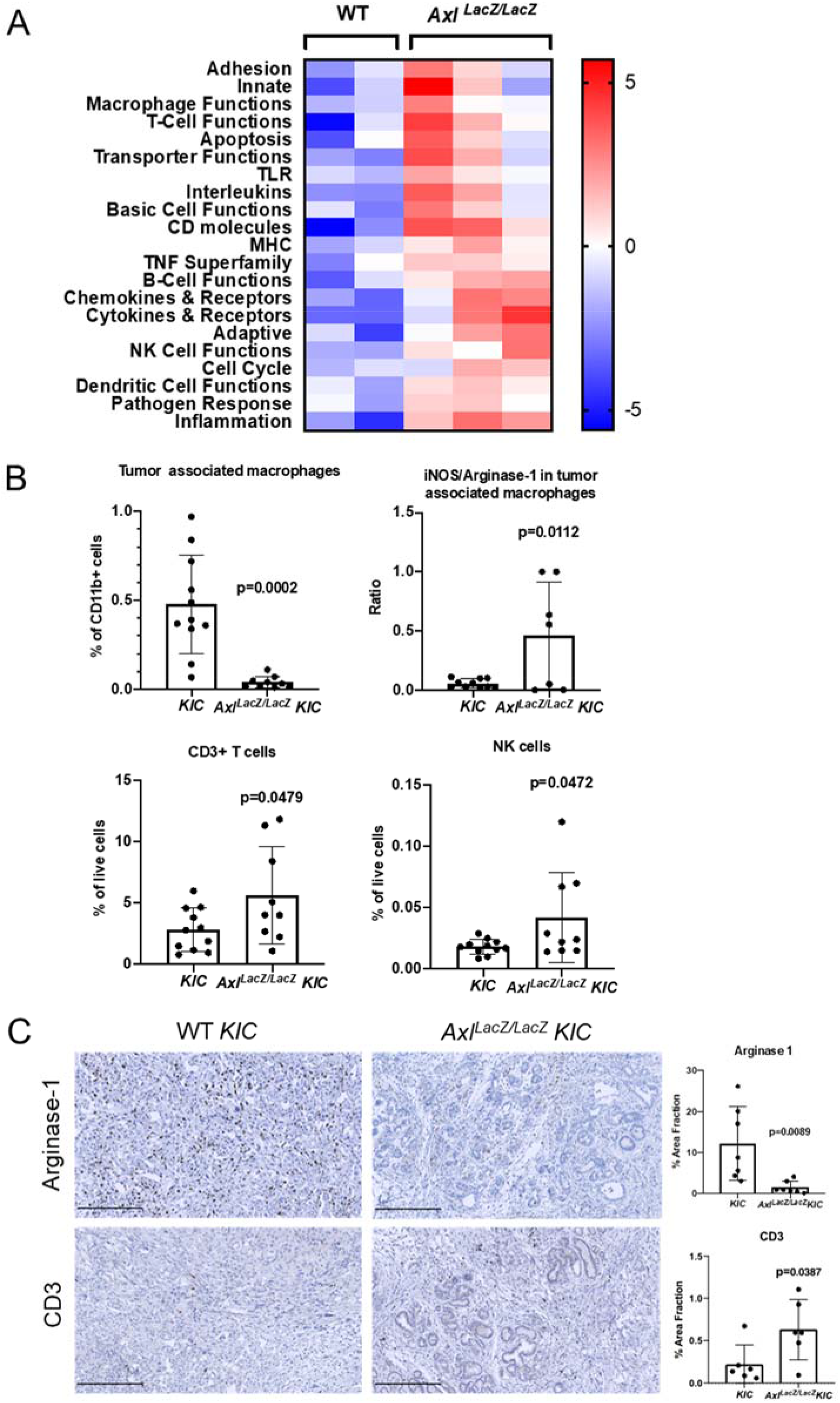
AXL deficiency results in a more active immune microenvironment. **A)** RNA was isolated from *KIC* (n=2) and *Axl*^*LacZ/LacZ*^ *KIC* (n=3) tumors and analyzed using Mouse PanCancer Immune Profiling Panel (NanoString Technologies). Upregulated (Red) and downregulated (Blue) gene programs/functions are shown. **B)** Flow cytometry of tumor-associated myeloid and lymphoid cells from *KIC* (n=11) and *Axl*^*LacZ/LacZ*^ *KIC* (n=9). Tumor associated macrophages (CD11b^+^ Ly6G^−^ Ly6C^−^ F4/80^+^ CD11c^+^ MHCII^+^), iNOS^+^ /Arginase-1^+^ macrophage ratio, CD3^+^ T cells, and CD335^+^ NK cells were analyzed. P values were generated by *t* test. **C)** Tumor tissues from *KIC* and *Axl*^*LacZ/LacZ*^ *KIC* mice were evaluated for indicated targets. Images were taken by Nanozoomer to cover the whole tissue and analyzed using ImageJ software. Quantification of % area fraction is shown. Each data point is the average of one sample. Data are displayed as mean ± SD. P values were generated by t-test. Scale bar, 250 μM.

To validate the Nanostring results, flow cytometry was performed on tumors from *KIC* (n=11) and *Axl*^*LacZ/LacZ*^ *KIC* (n=9). The number of total myeloid cell (CD11b^+^ cells) was not changed in *Axl*^*LacZ/LacZ*^ *KIC* (**Supplementary Fig. 4**). However, the number of tumor-associated macrophages was significantly reduced (**Fig. 3B**). In addition to decreased recruitment, iNOS/Arginase-1 ratio in these macrophages was up-regulated in *Axl*^*LacZ/LacZ*^ *KIC* (**Fig. 3B**), indicating a transformation from immunosuppressive macrophages to inflammatory macrophages. Consistent with this we identified dramatic decrease of Arginase-1 expression by immunohistochemistry in *Axl*^*LacZ/LacZ*^ *KIC* tumor tissue (**Fig. 3C**).

Flow cytometry also revealed an increased number of CD3^+^ T cells and CD335^+^ NK cells in *Axl*^*LacZ/LacZ*^ *KIC* while the CD4/CD3 and CD8/CD3 ratio were not changed (**Fig. 3B** and **Supplementary Fig. 4**). Elevated T cell infiltration (CD3) in *Axl*^*LacZ/LacZ*^ *KIC* was also observed by immunohistochemistry (**Fig. 3C**). These results suggest AXL is important for the immunosuppressive microenvironment of PDA and highlights that immune suppression might contribute directly to chemoresistance and metastasis (36, 37).

### AXL expression on tumor cells is critical for PDA progression and metastasis

We have demonstrated the effect of AXL depletion on different cell populations in PDA. To investigate which AXL-positive cell population contributes to PDA progression, AXL was ablated using CRISPR-Cas9 in PDA cells isolated from a *KPfC* tumor. In vitro functional assays demonstrated that loss of AXL (KO) does not influence cancer cell proliferation, but decreases cell migration and colony formation (**Supplementary Fig. 5**). Western blot analysis showed loss of AXL lead to decreased EMT transcription factor SLUG and mesenchymal marker Vimentin (**Fig. 4A**). Functional analysis showed that WT *KPfC* cells cultured in 30% matrigel+70% collagen (38) formed projections indicating invasiveness while AXL KO *KPfC* cells lost the ability to invade (**Fig. 4B**).

**Figure 4.**
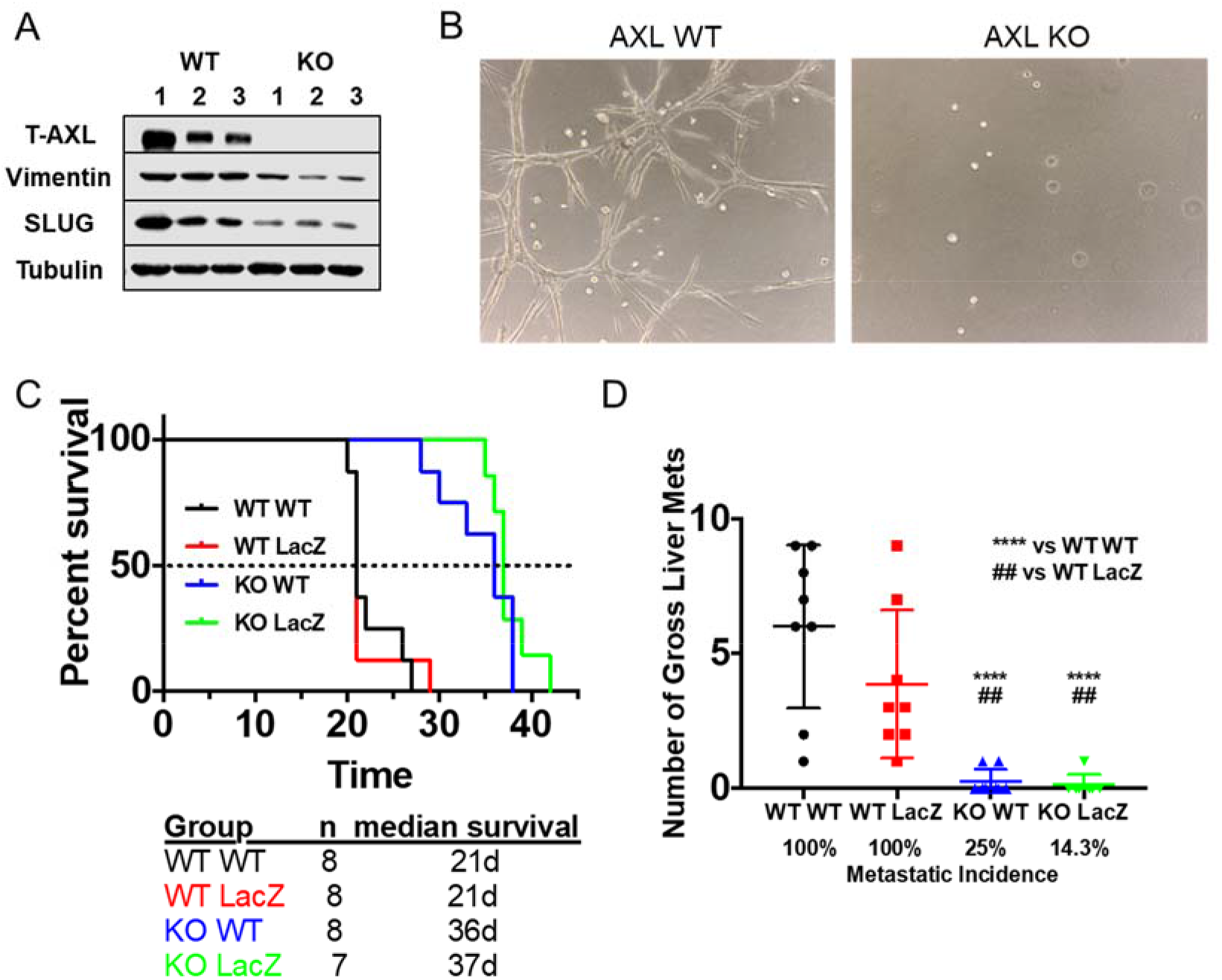
AXL expression on tumor cells is critical for PDA progression and metastasis. **A)** AXL was ablated out using CRISPR-Cas9 in *KPfC* tumor cells. Cell lysates from WT and AXL KO *KPfC* cells were evaluated for indicated targets by western blot. **B)** Representative pictures of WT and AXL KO *KPfC* cells grown in 30% matrigel+70% collagen for 4 days. **C)** WT and *Axl*^*LacZ/LacZ*^ C57BL/6 mice orthotopically injected with 250,000 cells/mouse WT and AXL KO *KPfC* cells were enrolled in a survival study. Groups were as follows: WT *KPfC* cells in WT C57BL/6 mice (n=8); WT *KPfC* cells in *Axl*^*LacZ/LacZ*^ C57BL/6 mice (n=8); AXL KO *KPfC* cells in WT C57BL/6 mice (n=8); AXL KO *KPfC* cells in *Axl*^*LacZ/LacZ*^ C57BL/6 mice (n=7). **D)** Number of gross liver metastases and metastatic incidence of the survival study in **(C)** are shown. ****P < 0.0001 vs WT *KPfC* cells in WT C57BL/6 mice; ## P < 0.01 vs WT *KPfC* cells in *Axl*^*LacZ/LacZ*^ C57BL/6 mice; by ANOVA.

To determine whether AXL expression on tumor cells or host cells is more critical to PDA progression, WT and AXL KO *KPfC* cells were orthotopically injected into WT and *Axl*^*LacZ/LacZ*^ C57BL/6 mice. Injection of AXL KO *KPfC* cells into WT mice (KO WT, blue, n=8, 36 days) resulted in longer median survival compared to WT mice injected with WT *KPfC* cells (WT WT, black, n=8, 21 days, p<0.0001, **Fig. 4C**). However, AXL-deficient mice injected with WT *KPfC* cells (WT LacZ, red, n=8, 21 days) showed the same median survival and metastases as WT mice injected with WT *KPfC* cells (p=0.9517, **Fig. 4D**). Similarly, AXL-deficient mice injected with AXL KO *KPfC* cell (KO LacZ, green, n=7, 37days) resulted in longer median survival compared to AXL-deficient mice injected with WT *KPfC* cells (p=0.0001, **Fig. 4C**). AXL-deficient mice injected with AXL KO *KPfC* cell showed similar median survival and metastases as WT mice injected with AXL KO *KPfC* cells (p=0.25, **Fig. 4C**). Gross liver metastases and metastatic incidence indicate that tumor cell expression of AXL was more important for metastasis than AXL expression by the host (**Fig. 4D**). Human PDA cell Panc1 and murine PDA cell Pan02 were also used to repeat the orthotopic experiment and the results were consistent (**Supplementary Fig. 6A** **and** **6C**). To investigate metastasis using Pan02 cells, splenic injection was performed in WT and *Axl*^*LacZ/LacZ*^ C57BL/6 mice and similar results were obtained (**Supplementary Fig. 6B**). These in vivo results indicate that AXL expression on tumor cells is critical for PDA progression and metastasis.

## Discussion

Multiple strategies targeting AXL for cancer therapy have been developed and are being investigated in the clinic (10, 39); however, AXL expression within PDA microenvironment and the contribution of different AXL-positive cell populations to PDA progression have not been investigated extensively. Our lab used scRNA-seq to identify AXL-positive cells and characterize the effect of global loss of AXL in PDA GEMMs. Our study demonstrates that AXL is expressed by tumor cells, fibroblasts and macrophages and AXL expression on tumor cells contributes significantly to PDA progression and sensitivity to chemotherapy. Our data also document that AXL supports the immune suppressive microenvironment of PDA.

The phenotype of *AXL-deficient* PDA GEMMs is consistent with results from Goyette et al. (40) who crossed *MMTV-Neu* mice with *AXL-deficient* mice and found a decrease in the frequency and the total number of lung metastatic lesions in the absence of AXL (40). The characterization of *AXL-deficient KIC* mice provides a potent rationale for use AXL as a therapeutic target, especially in combination with standard chemotherapy or potentially immune checkpoint blockade. In fact, pharmacological inhibition of AXL has similar effect in vivo. For example, bemcentinib (BGB324), a selective AXL inhibitor, sensitizes PDA mouse models to gemcitabine treatment and promotes an immune stimulatory landscape within the tumor (30). The efficacy of bemcentinib in combination with chemotherapy is being tested currently in pancreatic cancer patients (NCT03649321). Pharmacological inhibition of AXL has also been shown to decrease metastasis in breast cancer models (19, 41) and improve the efficacy of doxorubicin, BCR-ABL inhibitor, docetaxel, paclitaxel, EGFR inhibitor and nivolumab in breast cancer, chronic myelogenous leukemia, prostate cancer, uterine serous cancer, NSCLC and glioblastoma models, respectively (11, 19, 42–45).

The increased sensitivity of *AXL-deficient KIC* mice to gemcitabine is likely to due to a more differentiated phenotype tumor cell phenotype that includes the up-regulation of nucleoside transporters such as ENT1. Consistent with this, an EMT gene signature and AXL expression by cancer cells was associated with chemoresistance screens in multiple cancer types (4, 8). These observations fit with our scRNA-seq of PDA GEMMs that revealed *Axl* expression is elevated in advanced tumors and is associated with tumor cells that have a mesenchymal-like phenotype. Furthermore, *Axl* correlated negatively with classic epithelial markers. In fact, it was the most negatively associated marker among all the mesenchymal markers assessed. This highlights the use of AXL as a mesenchymal cancer cell marker. Our data also suggest that AXL-positive mesenchymal PDA cells are a major contributor to PDA progression and metastasis, further emphasizing the rationale to target this subpopulation of tumor cells. Recently, an anti-AXL chimeric antigen receptor T cell therapy has shown promising efficacy in vitro with myelogenous leukemia cells and triple negative breast cancer cells (46, 47). AXL has also been investigated as a target for antibody drug conjugates in melanoma, lung, pancreatic and cervical cancer models, some of which have advanced to phase 1 clinical testing (48). However, the mechanism of how AXL is up-regulated during cancer progression and how AXL maintains epithelial plasticity in this subpopulation of cancer cells remains to be elucidated. To this end, we have recently shown that AXL stimulates the activity of tank-binding kinase 1 (TBK1) to drive epithelial plasticity in PDA (49). Current studies are focused on understanding if this pathway is dependent on mutant KRAS signaling and how TBK1 stimulates EMT.

The immunosuppressive microenvironment in *KIC* tumor also contributes to chemoresistance and metastasis. We showed *AXL-deficient KIC* tumors have a more inflammatory microenvironment with an increase in T cell and NK cell recruitment and a potent decrease in tumor associated macrophages, which were also polarized from immunosuppressive (Arginase-1^+^) to anti-tumor (iNOS^+^) phenotype. This is consistent with the function of AXL in efferocytosis, where it has been shown as a stimulator of apoptotic cell phagocytosis by macrophages (50). The polarization and recruitment of immune cells can also be influenced by crosstalk between AXL-positive cancer cells and immune cells. The Shieh group (51) showed that conditioned media from AXL-positive oral squamous cell carcinoma cells polarized tumor-associated macrophages to an M2-like phenotype (51). Aguilera et al (52) also demonstrated that AXL on breast cancer cells suppresses MHC-I expression and promotes cytokine secretion that supports and recruits myeloid cells (52) further enhancing immune suppression. Tumor associated macrophages are critical regulators of cytotoxic lymphocytes (53). These immunosuppressive macrophages can also be induced by conditioned media from PDA cells and result in the secretion of deoxycytidine that competitively inhibits gemcitabine uptake and metabolism (54).

Using orthotopic implant models we show that AXL-positive PDA cells are critical to PDA progression and metastasis while AXL-positive stroma appears to be nonessential in PDA progression and metastasis. However, blockade of AXL on immune cells can lead to a reduction of metastasis in other types of cancers. Paolino et al. (25) reported that AXL, and its family members TYRO3 and MERTK, are ubiquitination substrates for E3 ubiquitin ligase casitas B-lineage lymphoma proto-oncogene-b in NK cells and inhibition of TAM receptors enhances the anti-metastatic activity of NK cells in breast cancer models (25). Indeed, pharmacological inhibition of AXL by BGB324 or RXDX-106 improves the efficacy of immune checkpoint blockade in glioblastoma and colon cancer (42, 55). Spatially specified excision of AXL will be required to determine the function of AXL on different populations of stroma cells in the tumor microenvironment.

In summary, we demonstrate that global loss of AXL results in improved outcome in a PDA GEMM and a more immune active microenvironment. Our studies also highlight that tumor cell expression of AXL is an important contributor to PDA progression and chemoresistance. Our results in aggregate further validate AXL as a therapeutic target in PDA.

## Materials and Methods

### Cell lines

Human pancreatic cancer cell line Panc-1 was obtained from ATCC, and murine pancreatic cancer cell line, Pan02, was obtained from the Developmental Therapeutics Program at the National Cancer Institute at Frederick, Maryland. *KPfC* cell lines were derived from *KPfC* (*Kras*^*LSL-G12D*^; *Trp53*^*fl/fl*^; *PDX*^*Cre*/+^) mice. All cell lines were cultured in DMEM (Invitrogen) containing 5% FBS and maintained in a humidified atmosphere with 5% CO_2_ at 37 °C. Human cell lines were DNA fingerprinted for provenance using the Power-Plex 1.2 kit (Promega) and confirmed to be the same as the DNA fingerprint library maintained by ATCC. All cell lines were confirmed to be free of mycoplasma (e-Myco kit, Boca Scientific) before use.

### Animal studies

*Kras*^*LSL-G12D*^; *Cdkn2a*^*fl/fl*^; *Ptf1a*^*Cre*/+^ (*KIC*) mice were generated as previously described (49). *Axl*^*LacZ/LacZ*^ mice were obtained from The Jackson Laboratory (*Axl*^*tm1Dgen*^, stock 005777) and were crossed into *KIC* resulting in *Axl*^*LacZ/LacZ*^ *KIC* on a C57Bl/6 background. At 45 days of age, male and female mice were randomized to receive vehicle (control, saline), or gemcitabine (25 mg/kg i.p. twice/week). Mice were weighed twice weekly and sacrificed when they were moribund or had >20%weight loss. Tissues were fixed in 10% formalin or snap-frozen in liquid nitrogen for further studies.

Male and female 6-week-old *C57BL/6* mice were obtained from an on-campus supplier. Male and female 6-week-old NON SCID mice were obtained from Charles River. *Axl*^*LacZ/LacZ*^ mice at the same age were generated in house. WT and AXL KO *KPfC* cells (2.5×10^5^), Pan02 cells (5×10^5^) and Panc1 cells (1 × 10^6^) were injected orthotopically (and splenically for Pan02). *KPfC* and Pan02 bearing mice were weighed twice weekly and sacrificed when they were moribund or had >20% weight loss. For splenically injection of Pan02, mice were sacrificed 7 weeks after injection. For Panc1 cells, mice were sacrificed 11 weeks after injection. Tissues were fixed in 10% formalin or snap-frozen in liquid nitrogen for further studies.

### Recombined *Cdkn2a* allele detection

Liver micrometastasis was assessed by quantitative reverse transcription PCR (RT-PCR) for the recombined *Cdkn2a(Ink4a/Arf)* allele. Briefly, frozen livers were homogenized in SDS lysis buffer (100 mM Tris, pH 8.8; 5 mM EDTA, 0.2% SDS, 100 mM NaCl) and digested at 56°C overnight. DNA was extracted using phenol/chloroform/isoamyl alcohol (25:24:1), and quantitative RT-PCR was performed using iTaq SYBR Green Supermix (Bio-Rad). The following validated primers were used for analysis of *Cdkn2a*: GCCGACATCTCTCTGACCTC (forward) and CTCGAACCAGGTTTCCATTG (reverse). Each sample was analyzed in triplicate.

### CRISPR-Cas9 knockout of AXL

Guide RNAs targeting *Axl* were cloned into the pX330 plasmid. The reconstructed plasmid was transfected into cells using lipofectamine 2000. 8 hours after transfection, medium was changed to normal medium. 48 hours after transfection, GFP^+^ cells were sorted out and plated in 96-well plates at 1 cell/well. AXL knockout cells were confirmed by Western blot. Sequences for guide RNAs are as follows: Human *AXL* GuideA for: caccgctgcctagccgaagctgat; rev: aaacatcagcttcggctaggcagc; Human *AXL* GuideB for: caccgcaccccttatcacatccgcg; rev: aaaccgcggatgtgataaggggtgc. Mouse *Axl* GuideA for: caccggagcttttccagccgaagc; rev: aaacgcttcggctggaaaagctcc; Mouse *Axl* GuideB for: caccgctcacaccccgtatcacatc; rev: aaacgatgtgatacggggtgtgagc.

### Nanostring analysis

*KIC* and *Axl*^*LacZ/LacZ*^ *KIC* tumor tissues were lysed in RLT lysis buffer and purified according to the manufacturer’s instructions (QIAGEN). RNA was sent to the Genomic and Microarray Core Facility at UTSW and analyzed using a preassembled nCounter PanCancer Immune Profiling Panel (mouse) and the nCounter system (NanoString Technologies) according to the manufacturer’s instructions. Samples were normalized based on the geometric means of the supplied positive controls and the panel of housekeeping genes, as recommended by the manufacturer. Only genes that were significantly different (P < 0.05; t test, false discover rate adjusted) and at least 1.5-fold differentially expressed between groups were considered.

### Bioinformatic analyses of single cell RNA sequencing data

Data were generated from our previous publication and analyzed as previously described (34). Spearman correlations of gene-gene expression in cancer cells were calculated using the correlation function from the stat package in R.

### Histology and tissue analysis

Formalin-fixed tissues were embedded in paraffin and cut in 5-mm sections. Sections were evaluated by H&E, Alcian blue and immunohistochemical analysis using antibodies specific for CD31 (Cell Signaling, 77699), KI67 (Abcam, ab15580), cleaved caspase-3 (CC3, Cell Signaling, 9661), αSMA (Biocare Medical CM 001), CDH11 (LifeSpan BioSciences, LS-B2308), E-cadherin (Cell Signaling, 3195), Vimentin (Cell Signaling, 5741), equilibrative nucleoside transporter 1 (ENT1, Abcam, ab135756), γH2AX (Novus, NB100-384), CD3 (Invitrogen, MA5-17043), Arginase 1 (Arg1, Cell Signaling, 93668), SOX9 (EMD Millipore, AB5535), PDGFRα (LifeSpan BioSciences, LS-B8655), and β-Galactosidase (Abcam, ab9361). Negative controls included omission of primary antibody. Color images were obtained with a Nikon Eclipse E600 microscope using a Niko Digital Dx1200me camera and ACT1 software, or Hamamatsu NanoZoomer 2.0-HT at Whole Brain Microscopy Facility of UTSW. Pictures were analyzed using ImageJ.

### MTS assay

The MTS colorimetric assay (Promega) was performed as per the manufacturer’s instructions. Cells were plated at 1,000 cells/well in tissue culture–treated 96-well plates. On each day, 20 μl/well MTS was added, followed by a 2-hour incubation at 37°C, and then the absorbance was read at 490 nm on a plate reader.

### Wound healing assay

Cells were cultured in 24-well tissue culture plates at high density (~90% confluence) in 0.5 ml of media with 5% FBS. Uniform scratches were made down the center of each well with a p200 pipette tip, cells were gently washed with PBS to remove the loose cell debris, and were cultured in media with 1% FBS. Images from the center of each well were taken at the indicated time points. The wound width was measured using ImageJ and percentage of wound healing (migration) was calculated based on the initial wound width.

### Liquid colony-forming assay

Cell lines were cultured in 6-well tissue culture plates at low density (200 cells/well) in 2 ml of media with 5% FBS and allowed to settle for 2 weeks or until marked colony formation. Cells were then fixed with 10% formalin and stained with crystal violet. Images were analyzed with ImageJ.

### Organotypic culture

For each cell line, 2000 cells were plated in 8-well chamber slides onto a base layer of growth factor-reduced Matrigel (5 mg/ml, BD Biosciences, Lot A6532) and collagen type I (1.5-2.1 mg/ml, BD Biosciences) and cultured for 2 days in a humidified 37°C incubator as previously described (38).

### Western blot analysis

*KPfC* cells were lysed using RIPA buffer (Cell Signaling, 9806), supernatants were recovered by centrifugation, protein concentration was measured using a Pierce BCA Protein Assay Kit (Thermo Fisher Scientific, 23225), and equal amounts of total protein were separated by SDS-PAGE. Proteins were transferred to nitrocellulose membranes (Bio-Rad), followed by a blocking in 5% BSA in TBST (0.05% Tween-20). The membranes were incubated overnight at 4°C with primary antibody, Actin (Sigma, A2066), AXL (Santa Cruz, sc-1096), SLUG (Cell Signaling, 9585), and Vimentin (Cell Signaling, 5741) followed by corresponding horseradish peroxidase-conjugated secondary antibody (Jackson ImmunoResearch). Specific bands were detected by using WesternSure PREMIUM chemiluminescent substrate (Li-Cor) on a Li-Cor imaging system (Odyssey-Fc).

### Flow cytometry

Tumors were digested with a cocktail containing collagenase I (45 μ/ml; Worthington), collagenase II (15 μ/ml; Worthington), collagenase III (45 μ/ml; Worthington), collagenase IV (45 μ/ml; Worthington), elastase (0.075 μ/ml; Worthington), hyaluronidase (30 μ/ml; MilliporeSigma), and DNase type I (25 μ/ml; MilliporeSigma) for 40 minutes at 37°C and passed through a 70-μm cell strainer (Falcon). Suspensions were washed twice with PBS and stained with Fixable Viability Dye (Thermo Fisher) for 1 hour. The cell suspensions were then washed and stained with antibodies detecting CD11b (BD Bioscience, 557657), Ly-6C (BD Bioscience, 562728), Ly-6G (BD Bioscience, 740953), F4/80 (Biolegend, 123132), CD11c (BD Bioscience, 564079), I-A/I-E (BD Bioscience, 562009), CD3 (BD Bioscience, 553061), CD4 (BD Bioscience, 562891), CD8 (BD Bioscience, 563332), and CD335 (Biolegend, 137621), for 1 hour at 4°C. Surface-stained cells were fixed, permeabilized, and stained for intracellular markers Arginase-1 (R&D Systems, IC5868P) and iNOS (Thermo Fisher, 17-5920-82). Cells were analyzed using FACS LSRFortessa SORP, and analysis was performed using FlowJo, with the help of the Moody Foundation flow cytometry facility at University of Texas Southwestern Medical Center.

### Statistical analysis

Data are reported as means ± SD. Statistical analyses of mouse model survival data were performed using analysis of variance (ANOVA) with Mantel-Cox test of significance difference using GraphPad Prism (version 8.0.1). Statistical analysis of immunohistorchemistry was performed with a 2-tailed t test or ANOVA using GraphPad Prism. For all analyses, P < 0.05 was considered statistically significant.

### Study approval

All animals were housed in a pathogen-free facility with 24-hour access to food and water. Experiments were approved by, and conducted in accordance with, the Institutional Animal Care and Use Committee at the University of Texas Southwestern Medical Center, which are compliant with the guidelines proposed by the NIH Guide for the Care and Use of Laboratory Animal.

## Author contributions

WD and RAB conceived and designed the study. WD, HH, ZW, JET, NB, YZ and NB acquired data and performed analysis and interpretation of data. WD wrote the manuscript. RAB reviewed and revised the manuscript. RAB supervised the study.

## Acknowledgements

We thank Drs. Jill Westcott, Gray Pearson, Srinivas Malladi and members of the Brekken lab for helpful discussion and Dave Primm for editorial assistance. The work was supported by NIH grants R01 CA192381 and U54 CA210181 Project 2 and the Effie Marie Cain Scholarship in Angiogenesis Research to RAB. The funders had no role in study design, data collection and analysis, decision to publish, or preparation of the manuscript.

---

**Supplementary Figure 1.**
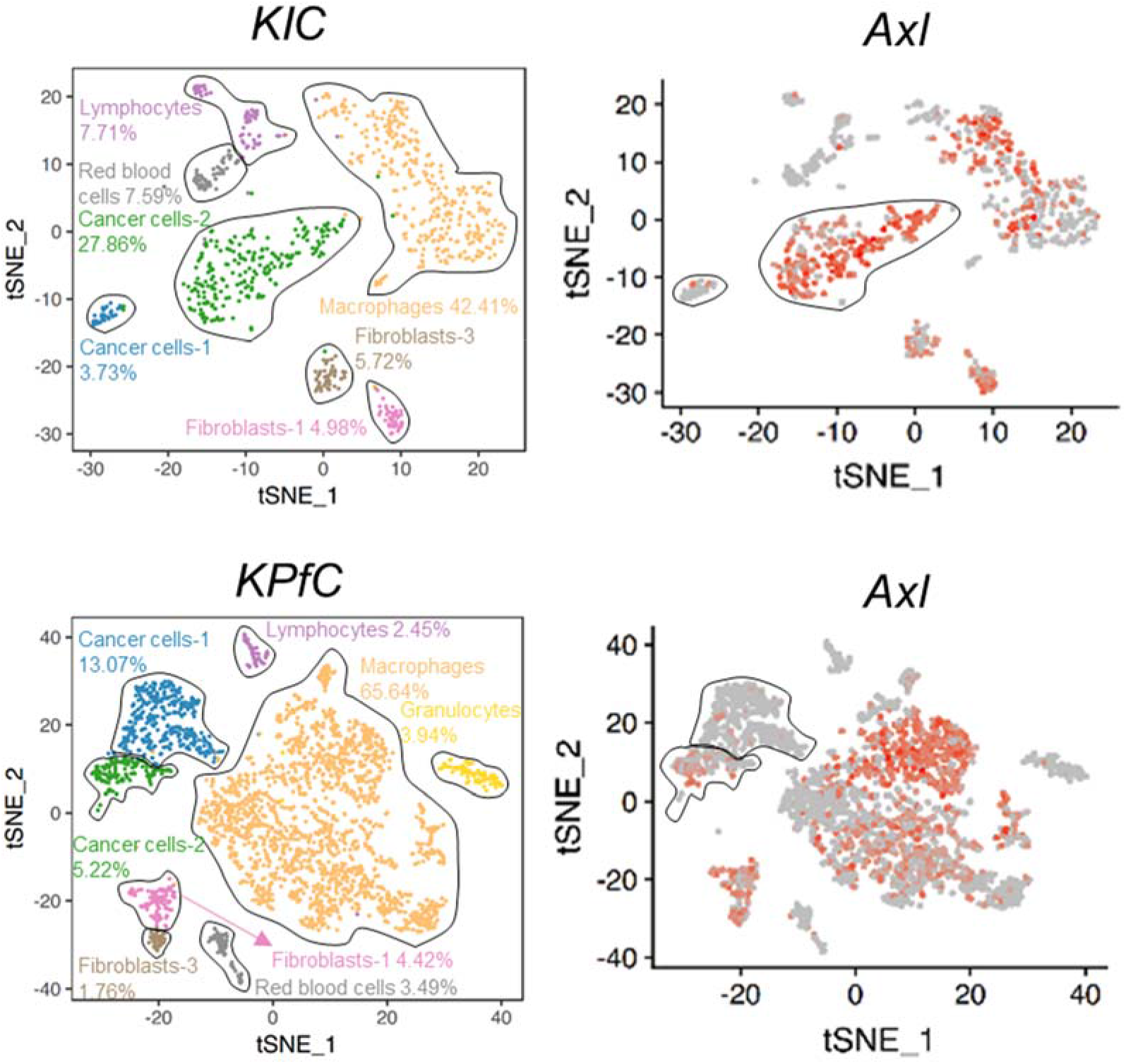
ScRNA-Seq was performed with tumors harvested from *KIC* or *KPfC* mice. tSNE plots of *Axl* expression for each cell population are shown. *Axl* expression is highlighted in red in the second tSNE plot for each model.

**Supplementary Figure 2.**
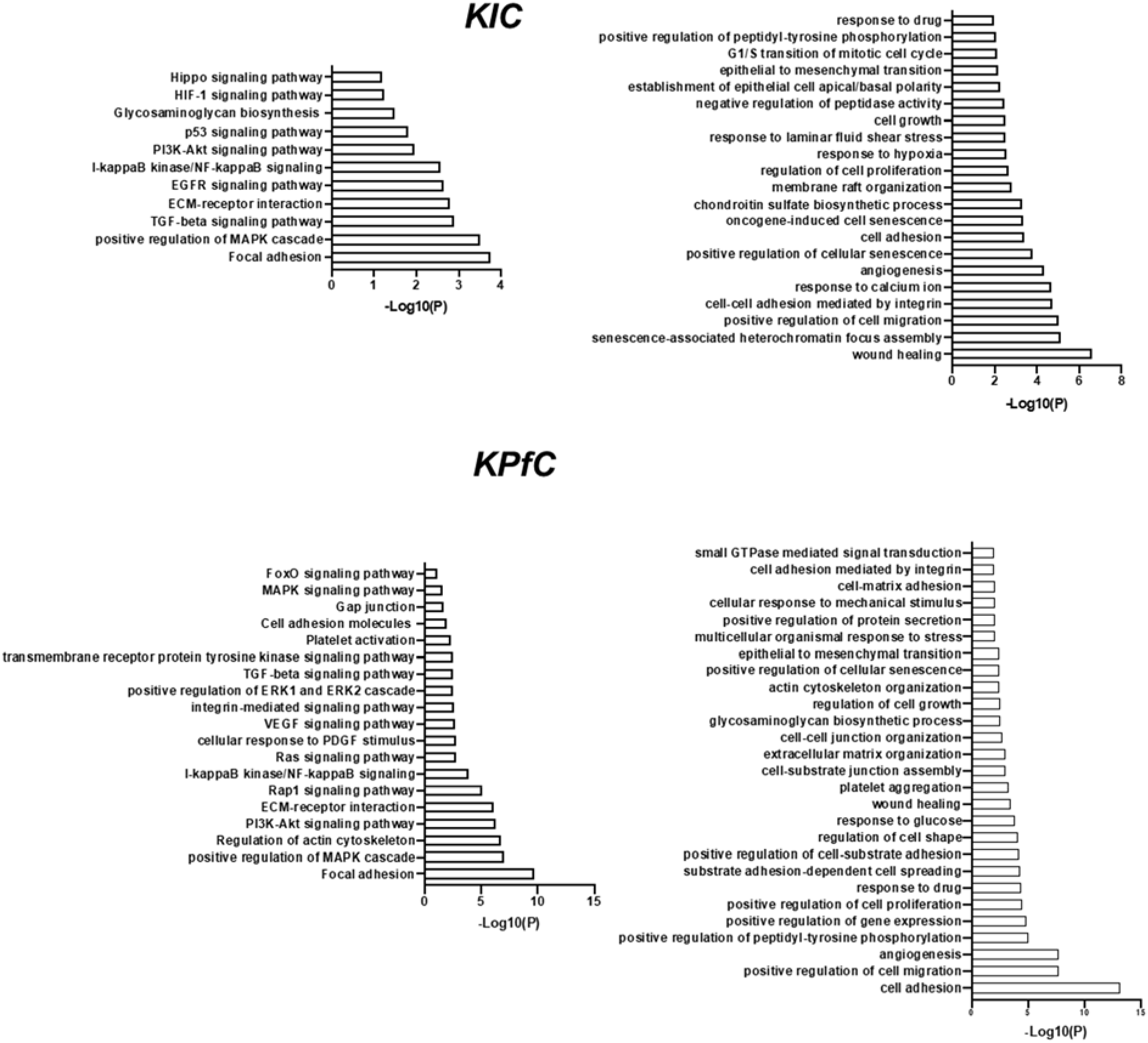
Gene Ontology (GO) analysis of gene clusters from AXL-positive cancer cell populations based on scRNA-Seq data.

**Supplementary Figure 3.**
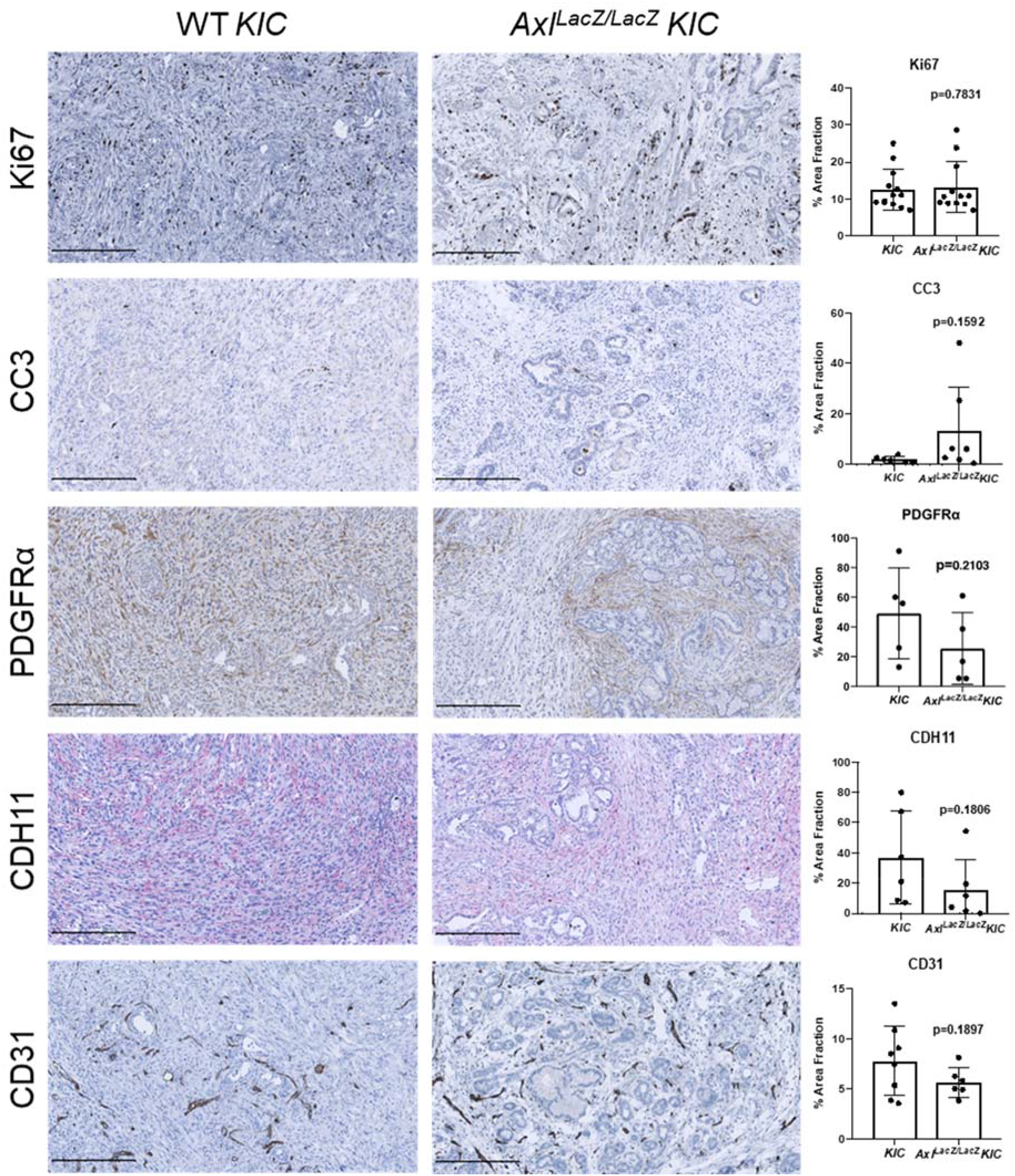
Tumor tissues from *KIC* and *Axl*^*LacZ/LacZ*^ *KIC* mice were evaluated for indicated targets. Images were taken by Nanozoomer to cover the whole tissue and analyzed using ImageJ software. quantification of % area fraction is shown. Each data point is the average of one sample. Data are displayed as mean ± SD. P values were generated by t-test. Scale bar, 250 μM.

**Supplementary Figure 4.**
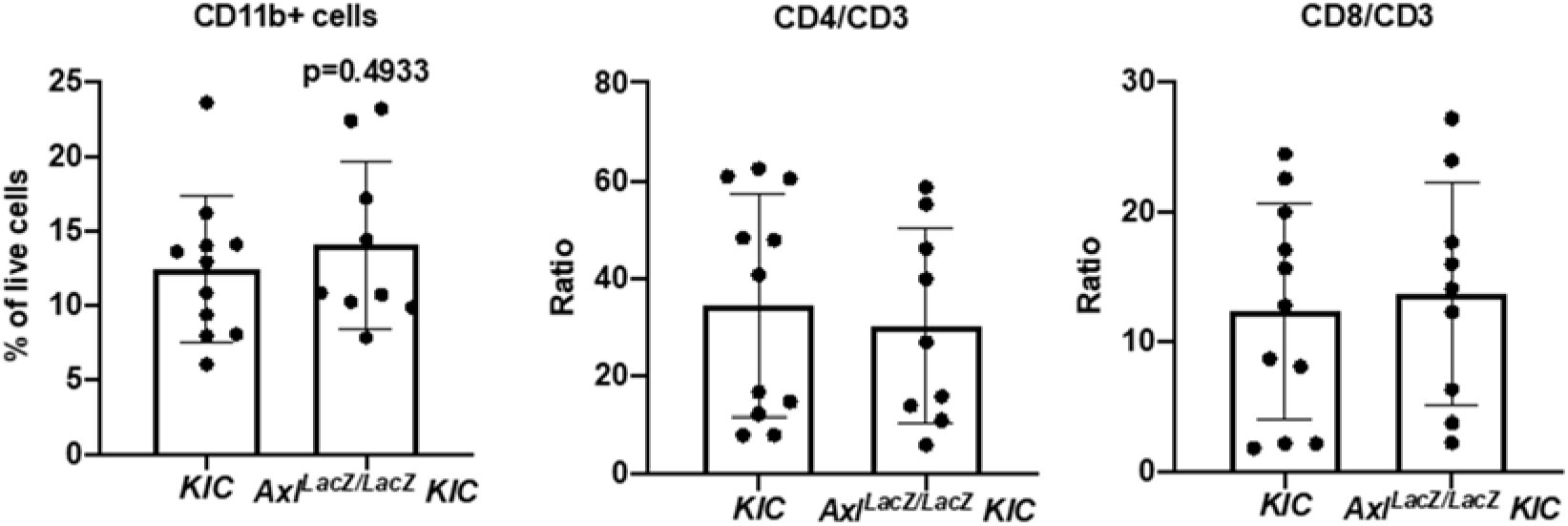
Flow cytometry of tumor-associated myeloid and lymphoid cells from *KIC* (n=11) and *Axl*^*LacZ/LacZ*^ *KIC* (n=9). CD11b^+^ myeloid cells, CD4/CD3, and CD8/CD3 ratio were analyzed. P values were generated by *t* test.

**Supplementary Figure 5.**
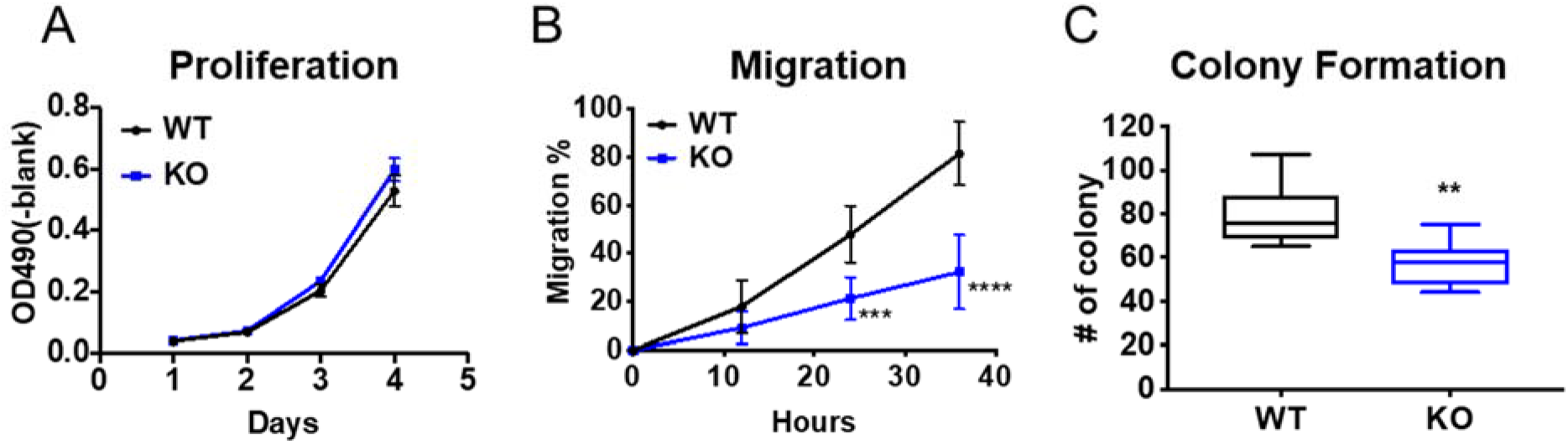
**A**) Cell proliferation was investigated in three different WT and AXL KO *KPfC* cells using MTS assay with eight technical replicates. **B)** Cell migration was investigated in three different WT and AXL KO *KPfC* cells by a “scratch” assay with three technical replicates. Monolayers of the indicated cells were wounded with a pipet tip. The cells were incubated in 1% serum media. Wound closure was monitored at 12, 24, and 36 hours and is reported as % wound closure. ***P < 0.001; ****P < 0.0001; by t-test. **C)** Colony formation (colonies/hpf) for three different WT and AXL KO *KPfC* cells with three technical replicates is shown by box plot (min to max). **P < 0.01; by t-test.

**Supplementary Figure 6.**
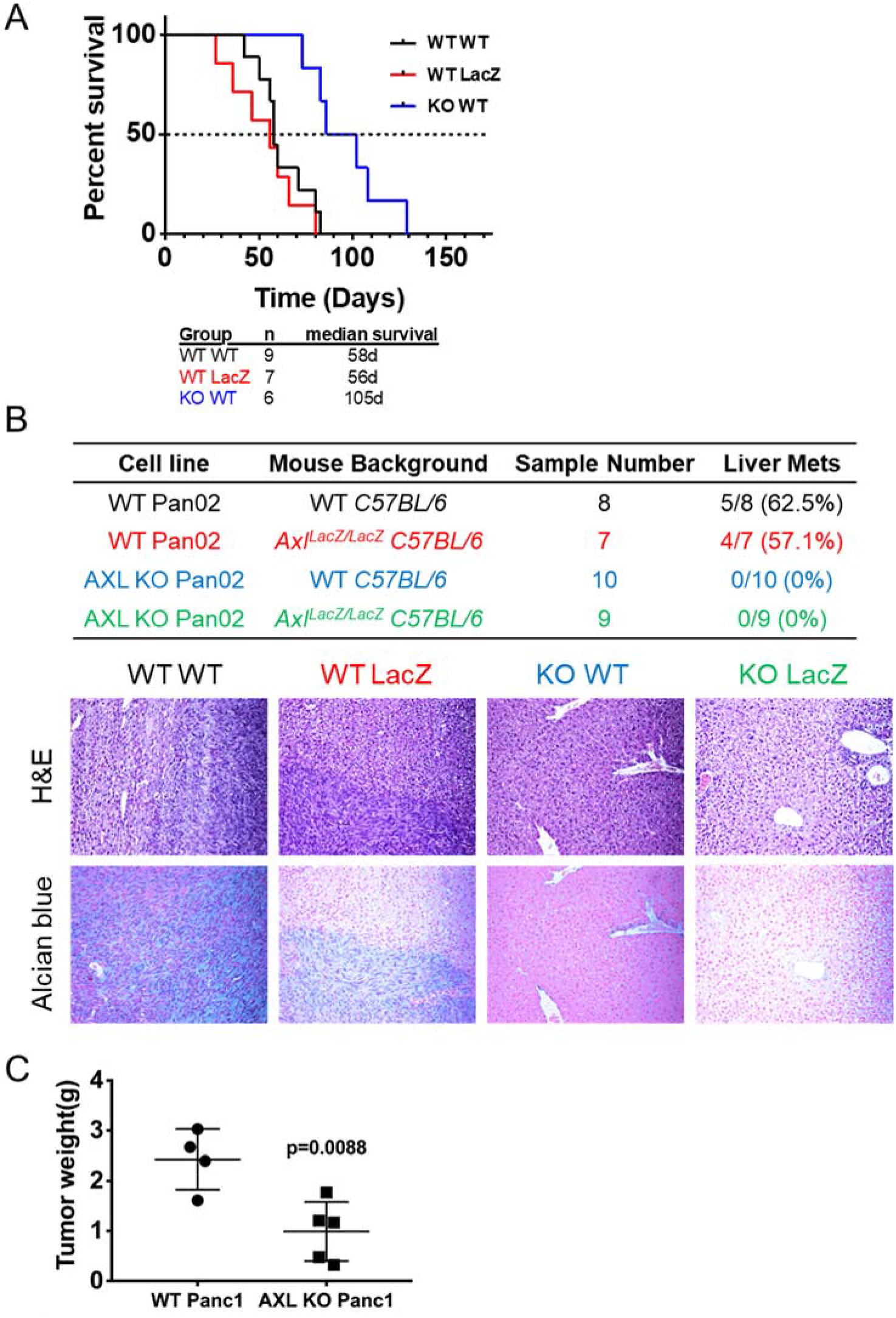
**A)** WT and *Axl*^*LacZ/LacZ*^ C57BL/6 mice orthotopically injected with 500,000 cells/mouse WT and AXL KO Pan02 cells were enrolled in a survival study. Groups were as follows: WT Pan02 cells in WT C57BL/6 mice (n=9); WT *KPfC* cells in *Axl*^*LacZ/LacZ*^ C57BL/6 mice (n=7); AXL KO *KPfC* cells in WT C57BL/6 mice (n=6). **B)** 500,000 cells/mouse WT and AXL KO Pan02 cells were splenically injected into WT and *Axl*^*LacZ/LacZ*^ C57BL/6 mice. Liver metastases were investigated 7 weeks after injection. Groups were as follows: WT Pan02 cells in WT C57BL/6 mice (n=8); WT *KPfC* cells in *Axl*^*LacZ/LacZ*^ C57BL/6 mice (n=7); AXL KO *KPfC* cells in WT C57BL/6 mice (n=10); AXL KO *KPfC* cells in *Axl*^*LacZ/LacZ*^ C57BL/6 mice (n=9). H&E and Alcian Blue stain were conducted in the liver tissues to confirm metastasis. Magnification, 10X. **C)** 1,000,000 cells/mouse WT and AXL KO Panc1 cells were orthotopically injected into NON SCID mice. Tumor weight were investigated 11 weeks after injection.

